# Asesino: a nucleus-forming phage that lacks PhuZ

**DOI:** 10.1101/2024.05.10.593592

**Authors:** Amy Prichard, Annika Sy, Justin Meyer, Elizabeth Villa, Joe Pogliano

## Abstract

As nucleus-forming phages become better characterized, understanding their unifying similarities and unique differences will help us understand how they occupy varied niches and infect diverse hosts. All identified nucleus-forming phages fall within the proposed Chimalliviridae family and share a core genome of 68 unique genes including chimallin, the major nuclear shell protein. A well-studied but non-essential protein encoded by many nucleus-forming phages is PhuZ, a tubulin homolog which aids in capsid migration, nucleus rotation, and nucleus positioning. One clade that represents 24% of all currently known chimalliviruses lacks a PhuZ homolog. Here we show that *Erwinia* phage Asesino, one member of this PhuZ-less clade, shares a common overall replication mechanism with other characterized nucleus-forming phages despite lacking PhuZ. We show that Asesino replicates via a phage nucleus that encloses phage DNA and partitions proteins in the nuclear compartment and cytoplasm in a manner similar to previously characterized nucleus-forming phages. Consistent with a lack of PhuZ, however, we did not observe active positioning or rotation of the phage nucleus within infected cells. These data show that some nucleus-forming phages have evolved to replicate efficiently without PhuZ, providing an example of a unique variation in the nucleus-based replication pathway.

## INTRODUCTION

The organization of cellular processes is essential for life. Depending on the species and context, this organization can be achieved by strategies such as subcellular compartmentalization, cytoskeletal structures, and phase separation. Viruses are no exception to this rule^1^. Nucleus-forming bacteriophages have been recently described as a group of bacterial viruses that form a proteinaceous replication compartment with many functional similarities to the eukaryotic nucleus^2,3^. The phage nucleus houses and protects phage DNA^2–6^; it is the site of DNA replication, recombination, and transcription^2,3,7^; and it exhibits selective import of proteins and export of mRNA^2,3,8,9^. In addition to the subcellular organization provided by the phage nucleus, some of these phages also assemble structures of mature virions known as bouquets^10^, and most currently characterized nucleus-forming phages encode a tubulin (PhuZ) that can perform diverse roles such as positioning the phage nucleus within the host cell, delivering capsids to the phage nucleus for genome packaging, and rotating the phage nucleus to distribute capsids evenly across its surface^2,3,11–17^. All currently characterized and putative nucleus-forming phages belong to one monophyletic clade that is proposed to be a viral family called Chimalliviridae^13^. This proposed family shares a core genome^13,18,19^ of 68 unique genes, including chimallin (ChmA) which makes up the phage nucleus shell^6,20,21^ and two multisubunit RNA polymerases (msRNAP)^7,22–24^. The two msRNAPs are the virion RNA polymerase (vRNAP), which is encapsidated along with the DNA, and the nonvirion RNA polymerase (nvRNAP), which localizes to the phage nucleus^22,25^. At the earliest stage of infection, the phage injects its DNA into the host cell with several proteins, including the vRNAP, that are required for early infection^25,26^, into a membrane-bound early infection intermediate^6,27–29^ called the EPI (early phage infection) vesicle^29^. Here, the vRNAP transcribes early genes. Later, the nvRNAP transcribes mid to late genes in the phage nucleus^7,24^.

Studying diverse chimalliviruses reveals novel insights into the variations in the nucleus-based phage replication mechanism. PhiKZ-like *Pseudomonas* phages PhiKZ^30^, 201phi2-1^31^, and PhiPA3^32^ were the first phages where the nucleus-based replication mechanism was characterized^2,3^. This intricate replication strategy was initially discovered while exploring the phage-encoded tubulin PhuZ, which makes triple-stranded filaments that position the viral replication compartment during 201phi2-1 infections^11,14,16^. Since then, many different variations of this replication strategy have been observed. For example, unlike the PhiKZ-like phages, *Serratia* phage PCH45 does not degrade the host chromosome^4,33^. *Erwinia* phage RAY also does not degrade the chromosome and makes a five-stranded tubulin filament^13^ rather than a three-stranded one^14^. *Escherichia* phage Goslar makes a vortex of PhuZ filaments rather than a bipolar spindle and centers its phage nucleus in a bulge in the host cell rather than at midcell^12^. However, one commonality between all of these phages is that they use tubulin-based PhuZ filaments to help organize their infections. Yet despite its high conservation, both dominant negative PhuZ mutants^2,3,11–13,15,16,34^ and genetic knockouts^35^ show that PhuZ is not essential for the replication of nucleus-based phages: The expression of dominant negative PhuZ only has a ∼50% decrease in infection efficiency^11^. This suggests that, while PhuZ is not essential for nucleus-based phage replication, it does improve infection efficacy, at least for the PhiKZ-like phages.

PhuZ is not only non-essential for the replication of nucleus-based phages, but it is also not part of the conserved core genome of the proposed Chimalliviridae family^13^. In fact, a recent publication has featured a nucleus-forming phage that does not encode a PhuZ homolog (*Vibrio* phage Ariel, also known as phiKT1028), much like its close relative *Vibrio* phage pTD1^36^. However, Ariel and pTD1 are part of a clade that includes PhuZ-encoding *Vibrio* phages Aphrodite1 and Eric (also known as phiKT1019)^36^.

In addition to a handful of phages like Ariel that lack PhuZ but belong to clades where most of the members encode a PhuZ homolog, an entire clade of chimalliviruses exists where none of the members have PhuZ homologs (Figure 1). This raises questions about how the organization and dynamics of chimallivirus infections without PhuZ differ from those with PhuZ: If PhuZ normally positions and rotates the phage nucleus in PhuZ-encoding chimalliviruses, do PhuZ-less phages use a different method for nucleus positioning and rotation, or are one or both of these functions dispensable? Do PhuZ-less phages still form a phage nucleus that reorganizes the host cell during infection but have evolved not to depend on a tubulin-based filament, or do they no longer make a phage nucleus, opting to use a different replication strategy from previously characterized chimalliviruses?

**Figure 1.**
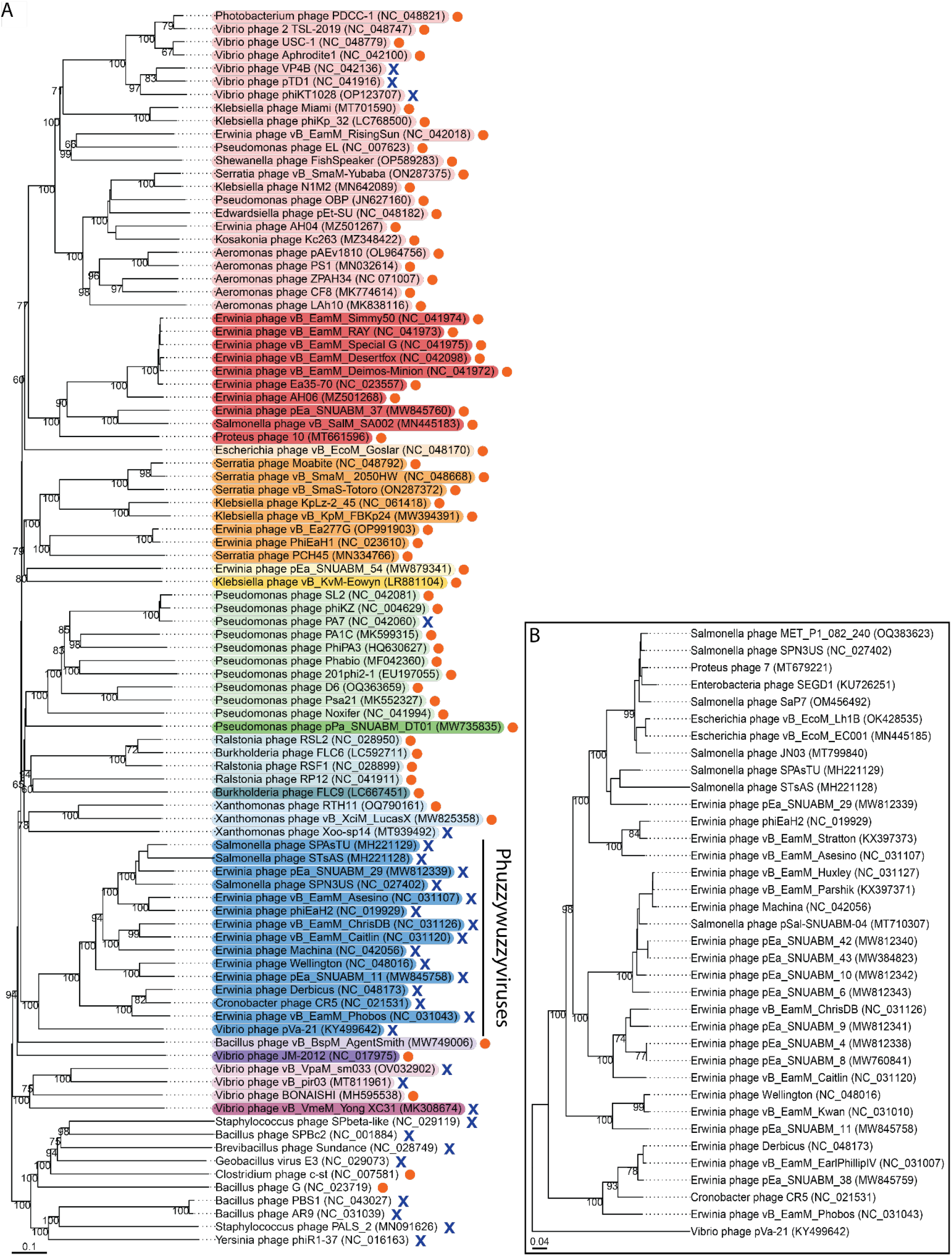
Phylogenetic trees of chimalliviruses. (A) A whole-genome phylogenetic tree made using VICTOR^45^ of the chimalliviruses, highlighted in rainbow colors according to predicted genus assignment (Figure S1), plus 10 outgroup representatives. Phages encoding a PhuZ homolog are marked with orange circles, and phages without PhuZ homologs are marked with blue X marks. The only clade where every member lacks PhuZ (besides singleton *Vibrio* phage Yong XC31) is highlighted in blue and labeled “Phuzzywuzzyviruses.” (B) A whole genome phylogenetic tree made using VICTOR^45^ of all currently known members of the phuzzywuzzyvirus clade. For both trees, bootstrap values are displayed on the branches.

In order to address these questions, we studied the PhuZ-less chimallivirus vB_EamM_Asesino (here termed simply Asesino). Asesino belongs to the clade of chimalliviruses where all members lack PhuZ (Figure 1A,B). The host of Asesino is *Erwinia amylovora*, an agriculturally important plant pathogen that infects plants of the Rosaceae family, including apple and pear trees^37^. *E. amylovora* is a good model for studying the variations in nucleus-based phage replication since many diverse chimalliviruses across five different clades have been isolated that infect this host^38–40^ (Figure 1A). By studying Asesino, we set out to determine how nucleus-based phage replication in the absence of a PhuZ homolog compares to previously characterized nucleus-forming phages.

## RESULTS

### Phylogenetics

Analyzing the conservation of PhuZ across Chimalliviridae members reveals a distinct clade within the proposed family where every member lacks PhuZ (Figure 1A, blue). All known members of this clade are shown in Figure 1B. Twenty-three of these phage infect *Erwinia*, and the remaining thirteen phage infect *Salmonella, Proteus, Escherichia, Enterobacter, Vibrio*, or *Cronobacter* (Figure 1B). Based on the branch lengths in the whole genome tree, these phage appear to be descended from a common ancestor that lost its PhuZ homolog and subsequently evolved to infect a range of bacteria (Figure 1A). *Erwinia* phage Asesino is part of this clade of PhuZ-less phages, hereafter referred to as phuzzywuzzyviruses due to their apparent evolutionary loss of PhuZ. In contrast to phuzzywuzzyvirus Asesino, previously characterized *Vibrio* phage Ariel (also known as phiKT1028) groups with a different clade of chimalliviruses, the *Tidunavirus* genus, and appears to have lost its PhuZ homolog more recently since it has closer relatives, such as *Vibrio* phage Aphrodite1, that encode PhuZ (Figure 1A, light red). The multiple instances of nucleus-forming phages that appear to have lost PhuZ (Figure 1A, blue X marks), including the puzzywuzzyvirus and *Tidunavirus* clades, suggest that it is adaptive to lose PhuZ in some circumstances.

### Asesino phage nucleus formation

To determine if Asesino forms a phage nucleus, we infected *E. amylovora* with Asesino, stained the DNA with DAPI, and imaged the infected cells at 75 minutes post-infection (mpi) using fluorescence microscopy. We observed a bright central zone of concentrated DNA in infected cells consistent with formation of a phage nucleus. (Figure 2A, magenta). To confirm that this concentrated mass of DNA is a phage nucleus made by Asesino, we tagged its ChmA homolog (gp039) with GFP on the N-terminus and expressed it during infection. GFP-ChmA formed a ring enclosing the bright DAPI-stained region, confirming the presence of a chimallin-based phage nucleus (Figure 2B, cyan). Similar to previously characterized *Erwinia* phage RAY^13^ and *Serratia* phage PCH45^4^, Asesino does not noticeably degrade the host DNA, leaving some DAPI staining outside the phage nucleus. This extranuclear DAPI staining co-localizes with GFP-tagged histone-like protein H-NS, which coats bacterial DNA, further suggesting that it is host DNA (Figure 2C).

**Figure 2.**
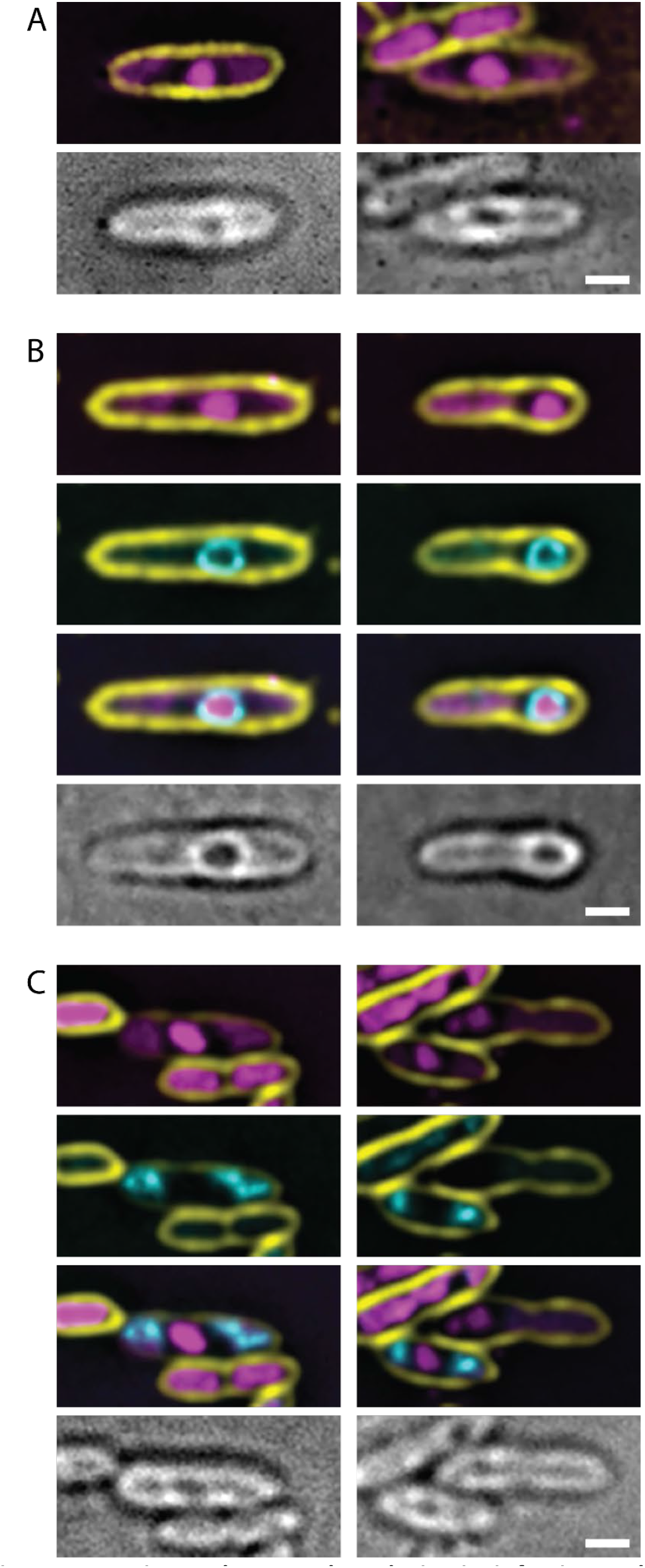
Asesino makes a nucleus during its infection cycle. (A) DAPI staining shows a bright circular region consistent with a phage nucleus, as well as extranuclear DNA consistent with undegraded bacterial DNA. (B) Asesino’s chimallin homolog (gp039) tagged with GFP encloses the bright DAPI signal. (C) *E. amylovora* H-NS tagged with GFP localizes with the extranuclear DNA during infection, supporting the idea that this is undegraded host DNA. For all panels, yellow is FM4-64 (membrane stain), magenta is DAPI (DNA stain), cyan is GFP, and grayscale is brightfield. Scale bars represent 1μm.

### Infection organization

We next assessed whether Asesino organizes its infection similarly to previously characterized nucleus-forming phages. The phage-encoded metabolic enzyme thymidylate kinase (gp117) which is predicted to contribute to the synthesis of nucleotide pools localizes to the cytoplasm (Figure 3A), whereas the nvRNAP β’ subunit 1 (gp040), which is required for transcription of mid to late phage genes^7^, and the phage-encoded RecA-like recombination protein (UvsX, gp238) localize to the phage nucleus (Figure 3B,C). Virion structural proteins (major capsid protein gp091 and tail protein gp166) localize between the phage nucleus and the bacterial nucleoids (Figure 3D,E), suggesting that mature viral particles accumulate in the space between the phage nucleus and the host chromosome. These results are similar to *Erwinia* phage RAY, which forms a phage nucleus but does not appear to assemble its mature viral particles into bouquets^13^.

**Figure 3.**
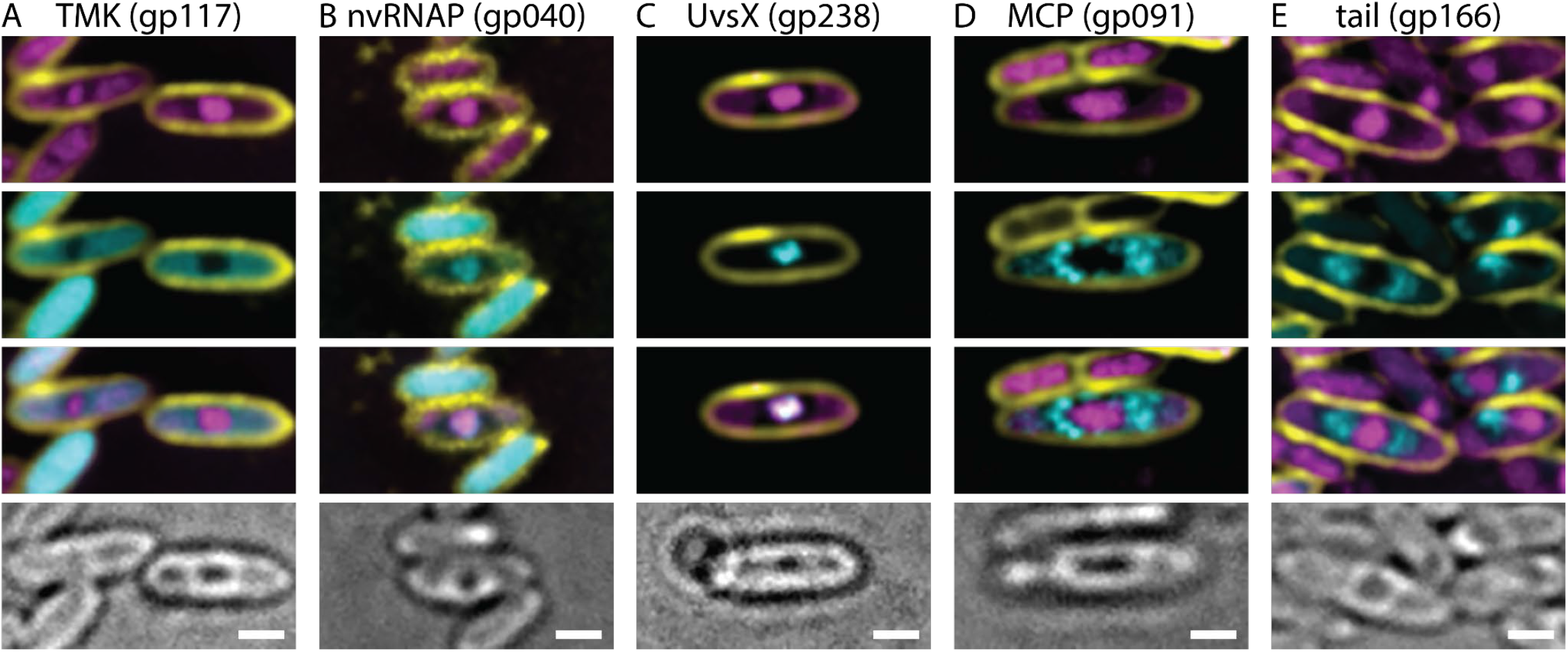
Protein localization in Asesino is consistent with other nucleus-forming phages. (A) Thymidylate kinase is localized in the cytoplasm. (B) A nonvirion RNA polymerase subunit and (C) RecA homolog UvsX are localized in the nucleus. (D) The major capsid protein and (E) a tail protein localize around the phage nucleus. For all panels, yellow is FM4-64 (membrane stain), magenta is DAPI (DNA stain), cyan is GFP, and grayscale is brightfield. Scale bars represent 1μm.

### Phage nucleus dynamics in the absence of PhuZ

Since Asesino makes a phage nucleus, we next determined if the effects we normally attribute to PhuZ were indeed missing during its replication cycle, or if they were somehow supplemented by other genes. To do this, we analyzed nucleus rotation dynamics and positioning. When measured at 60 mpi, nucleus rotation was not observed in time-lapse microscopy (n=46), supporting the idea that PhuZ is necessary for phage nucleus rotation.

Asesino phage nucleus positioning was quantified and compared to *Erwinia* phage RAY since both phages infect the same host, and neither degrades the host genome. In RAY, it was shown that lack of host genome degradation influences phage nucleus positioning, likely because the bacterial nucleoids take up space and occlude regions of the cell near the poles, preventing the phage nucleus from reaching those spaces even when PhuZ activity was inhibited using a dominant negative mutant^13^. Asesino appears not to position its nucleus as actively as RAY: When only one bacterial nucleoid is present, Asesino nucleus positioning is not biased toward midcell (Figure 4A). However, similar to RAY, it appears that the presence of bacterial nucleoids aid in positioning Asesino’s nucleus closer to midcell when there is one on each side (Figure 4B).

**Figure 4.**
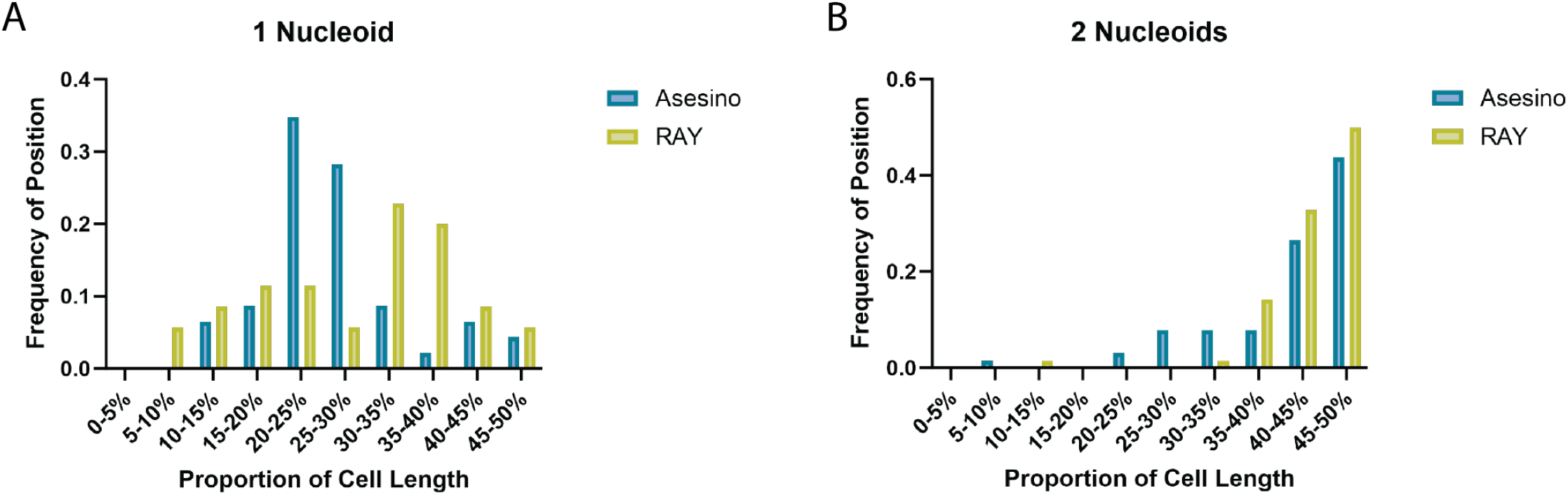
Phage nucleus positioning in Asesino compared to RAY. (A) When only one bacterial nucleoid is present, the Asesino phage nucleus has a bias away from midcell positioning, with most phage nuclei being found between the pole and midcell (20-30% of the cell length, blue), but the RAY phage nucleus has a slight bias toward midcell positioning (light green). (B) When two bacterial nucleoids are present, both the Asesino (blue) and RAY (light green) phage nuclei are positioned at or near midcell, with Asesino having a slightly wider distribution.

## DISCUSSION

Even before the discovery of the phage nucleus, PhuZ had been shown to perform an important role in positioning phage DNA during the infection cycle of nucleus-forming phages^11,14,16^. Yet upon further investigation, it was discovered to be an accessory factor that is not essential for the nucleus-based phage replication cycle^11,13,34,35^. In studies using dominant negative or null mutants of PhuZ, the phage nucleus becomes lodged adjacent to the cell pole, occluding a significant fraction of the surface area available for capsid docking and DNA packaging which is proposed to be the reason for decreased virion production^11,35,41^. How, then, do the many chimalliviruses that lack a tubulin homolog entirely (Figure 1A,B) avoid the problem of large portions of the phage nucleus surface area becoming inaccessible by apposition to the membrane? Here we studied the *Erwinia*-infecting phuzzywuzzyvirus Asesino, in order to gain a better understanding of PhuZ-less chimallivirus replication. We found that Asesino forms a nucleus-like structure that compartmentalizes phage DNA away from the cytoplasm and partitions proteins into two distinct compartments, similar to other nucleus-forming phages^2–4,12,13,36^. During Asesino infection, the bacterial chromosome is not degraded. This contrasts with *Vibrio* phage Ariel, the only other PhuZ-less phage whose replication mechanism has been studied to date, in which phage replication is accompanied by the degradation of bacterial DNA and the swelling of host cells into spheroids^36^. Cell swelling and bacterial genome degradation could be a mechanism for increasing diffusion around the phage nucleus. However, our results suggest that Asesino does not cause such drastic changes in cell shape, nor does it noticeably degrade the host DNA (Figure 2). The Asesino phage nucleus is preferentially localized near midcell when there are two bacterial nucleoids present (Figure 4B), and near the one-quarter cell position when only one nucleoid is present (Figure 4A). This is similar to RAY phage nucleus positioning when a dominant negative PhuZ mutant is expressed^13^. By leaving the host DNA intact, the Asesino phage nucleus competes for space with bacterial nucleoids, which appear to act as obstacles that help center the Asesino nucleus when two nucleoids are present (Figure 4B), as has been proposed for *Erwinia* phage RAY^13^. Additionally, it has been proposed that polysomes can be involved in physically separating DNA in bacterial cells^42,43^. A similar mechanism could be used by Asesino, where actively translating ribosomes cluster around the phage nucleus and act as physical buffers that center it between the host nucleoids.

We propose that each phage has evolved its own unique proteins and mechanisms for optimizing phage production and specializing on the microenvironment within the bacterial strain it infects. For instance, PhiKZ-like phages inject their genomes at the cell pole and cause the *Pseudomonas* cell to swell at midcell^2,3^ (Figure 5A). This allows the phage nucleus to grow larger, replicate more phage genomes, and therefore produce more phage particles. In these phages, the PhuZ spindle positions the phage nucleus at midcell, where the bulge forms. In the absence of a tubulin spindle, the phage nucleus forms and grows at the cell pole, where a large fraction of its surface would be occluded, limiting exchange of molecules into and out of the phage nucleus and restricting the surface area accessible for capsid docking, as happens for PhiKZ-like phages with dominant negative or null mutants of PhuZ^2,11,15,34,35^ (Figure 5B). It is tempting to speculate that centering the phage nucleus and bulging at midcell co-evolved to optimize progeny production in these phages. In the case of the PhuZ-less phage Ariel, it has adapted to its own unique challenge of replicating in *Vibrio parahaemolyticus* without a PhuZ spindle by degrading the host chromosome and causing the cell to swell and become spheroidal^36^ (Figure 5C). A cytoskeletal filament would not be required to find the middle of an isotropic cell, thus alleviating the selective pressure for positioning: It is impossible for the phage nucleus to get lodged in a low-diffusion environment such as the cell pole when no cell pole exists. In the case of *Erwinia* phage Asesino, the host cell remains anisotropic, does not exhibit significant cell swelling, and does not appear to contain a cellular region that would be more conducive for phage replication, although it does appear to leave the host DNA intact (Figure 5D), which could aid in positioning its phage nucleus^13^ (Figure 4), as discussed above. Thus, Asesino may have evolved these traits to benefit its replication efficiency and best take advantage of its host in response to selection pressures that we currently do not understand.

**Figure 5.**
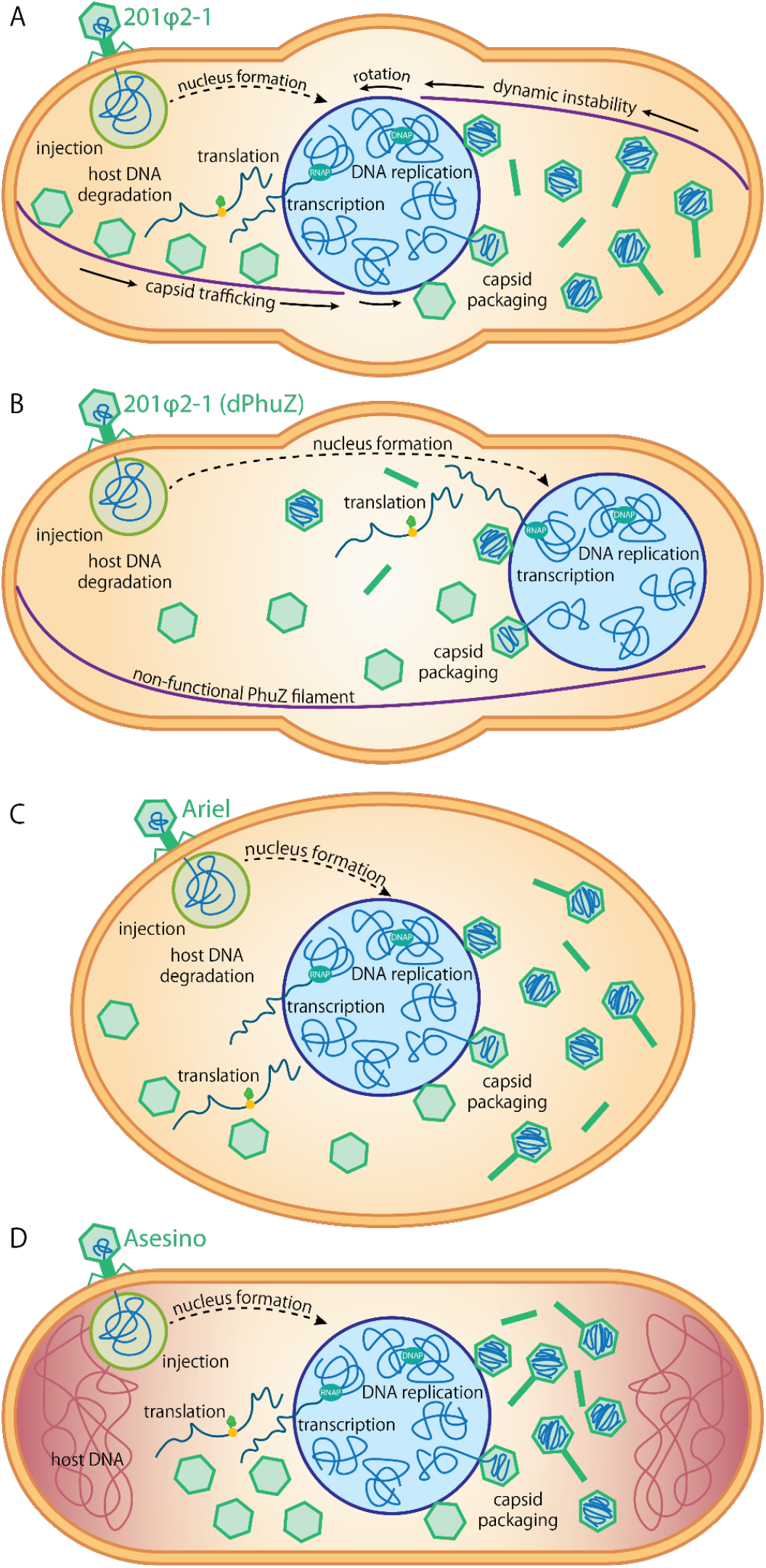
Infection Models. Diagrams comparing (A) a PhiKZ-like *Pseudomonas* phage 201phi2-1 infected cell^2^; (B) a 201phi2-1 infected cell expressing a catalytically dead, dominant negative PhuZ mutant^2^; (C) a *Vibrio* phage Ariel infected cell^36^; and (D) an *Erwinia* phage Asesino infected cell.

While the phage nucleus provides protection against DNA-targeting host defenses by physically excluding harmful nucleases^4,5,8^, nucleus-forming phages are not exempt from arms races with their hosts^4,33,35,44^ and are under constant selective pressure from other types of host defense systems, including RNA-targeting CRISPR systems which target conserved genes. Since phage are incredibly adaptable and evolve quickly, it is not surprising to find clades of phage where non-essential conserved genes such as PhuZ have been lost. Depending on the circumstances, loss of PhuZ may be advantageous (e.g. to avoid Cas13 targeting) or disadvantageous (e.g. random loss through genetic drift that slightly decreases replication efficiency). Once lost, it may be more difficult for nucleus-forming phages to reacquire genes, compared to canonically-replicating phages, since the phage nucleus can act as a barrier that limits genetic exchange^34^.

While all chimalliviruses characterized to date share the nucleus-based phage replication strategy, each has its own unique variation on this theme. PhuZ-encoding chimalliviruses, such as the PhiKZ-like phages, as well as PhuZ-less chimalliviruses, such as *Vibrio* phage Ariel and *Erwinia* phage Asesino, all use unique methods for ensuring efficacious phage replication in their respective hosts (Figure 5). While each strategy employed by chimalliviruses allows them to specialize on their specific host in their own unique ways, it is remarkable that at the same time, they share the core conserved nucleus-based phage replication mechanism.

## METHODS

### Phylogenetic Analyses

Whole genome phylogenetic trees were made by running VICTOR on amino acid data using the D6 formula^45^ and annotated in Illustrator (Adobe) based on VICTOR annotations and the presence of PhuZ based on BLAST results.

### Bacterial Growth and Phage Propagation Conditions

*Erwinia amylovora* ATCC 29780 was generally grown on LB plates at room temperature or in rotating liquid LB cultures at 30°C. *Erwinia* phage Asesino (vB_EamM_Asesino) and *Erwinia* phage RAY (vB_EamM_RAY) were propagated by infecting 0.5mL of dense *E. amylovora* from an overnight culture grown in NBSYE (13g/L Difco Nutrient Broth, 2.5g/L Bacto-Yeast Extract, 5g/L sucrose) with 10μL phage. After waiting 15 minutes for the phage to adsorb, the infected cells were mixed with 4.5mL molten (∼55°C) NBSYE 0.5% top agar and poured over an NBSYE 1.6% agar plate. These plates were incubated overnight at room temperature. The following day, fresh phage lysates were collected by soaking the plates with 5mL phage buffer (10mM Tris pH 7.5, 10mM MgSO_4_, 4g/L NaCl, 1mM CaCl_2_) for 5 hours, collecting the liquid by aspiration, pelleting the cell debris by centrifugation (3220 rcf for 10 min), and filtering the clarified phage lysate through a 0.45μm filter. Phage lysates were stored at 4°C.

### Fluorescence Microscopy

Fluorescently labeled phage proteins were tagged with GFPmut1 and expressed from the pHERD-30T plasmid under the inducible control of the AraBAD promoter. 15μg/mL gentamicin sulfate was used for selection prior to microscopy. To prepare bacterial cells for microscopy, imaging pads were prepared (1% agarose, 25% LB, and 0-0.2% arabinose depending on the particular construct imaged) in welled slides and inoculated with *E. amylovora*. The slides were incubated at 30°C in humid chambers for 3 hours for the bacteria to begin actively growing. They were then moved to room temperature and infected with 10μL undiluted phage lysate. 8μL of dye mix (12.5μg/mL FM4-64, 25μg/mL DAPI, 25% LB) was added to the microscopy pads immediately prior to imaging. Slides were imaged using a DeltaVision Elite deconvolution microscope (Applied Precision) and deconvolved using the aggressive algorithm in the DeltaVision softWoRx program (Applied Precision). All analysis was performed prior to deconvolution.

### Nucleus Rotation Analysis

Time lapses (2 minutes long with 4 second intervals) were obtained between 60-90 mpi and observed to detect rotation. Since no rotating phage nuclei were observed, no further analysis was performed to determine rotation dynamics.

### Phage Nucleus Positioning

The length of the cell and the distance between the center of the phage nucleus and the cell pole was measured using FIJI. The position of the nucleus within the cell was calculated as the ratio of the distance from the phage nucleus to the pole over the length of the cell. The data were graphed as a relative frequency histogram using Prism (GraphPad).

## ACKNOWLEDGEMENTS

We thank Julianne Grose from Brigham Young University for graciously sharing *Erwinia* phages Asesino and RAY with us and for offering helpful comments on the manuscript. We also thank Kevin Corbett, Ann Aindow, and Emily Armbruster for their helpful comments and suggestions on the manuscript. This work was supported by an Emerging Pathogens Initiative grant from the Howard Hughes Medical Institute (to J.P., E.V., and J.M.) and the National Institutes of Health R01-GM129245 (to J.P. and E.V.).

## AUTHOR CONTRIBUTIONS

Conceptualization, A.P., A.S., and J.P.; methodology, A.P.; validation, A.P. and A.S.; formal analysis, A.P.; investigation, A.P. and A.S.; resources, and J.P.; data curation, J.P.; writing - original draft preparation, A.P. and J.P.; writing - review and editing, A.P., A.S., J.M., E.V., and J.P.; visualization, A.P.; supervision, J.M. and J.P.; project administration, J.P.; funding acquisition, J.P., E.V., J.M.

## DATA AVAILABILITY

All raw microscopy images for analysis and publication in this paper are deposited in a Mendeley dataset (DOI: 10.17632/5tpmyz66sz.1).

