## Supplementary figures and images for "Asesino: a nucleus-forming phage that lacks PhuZ"

### Supplemantary Figure S1

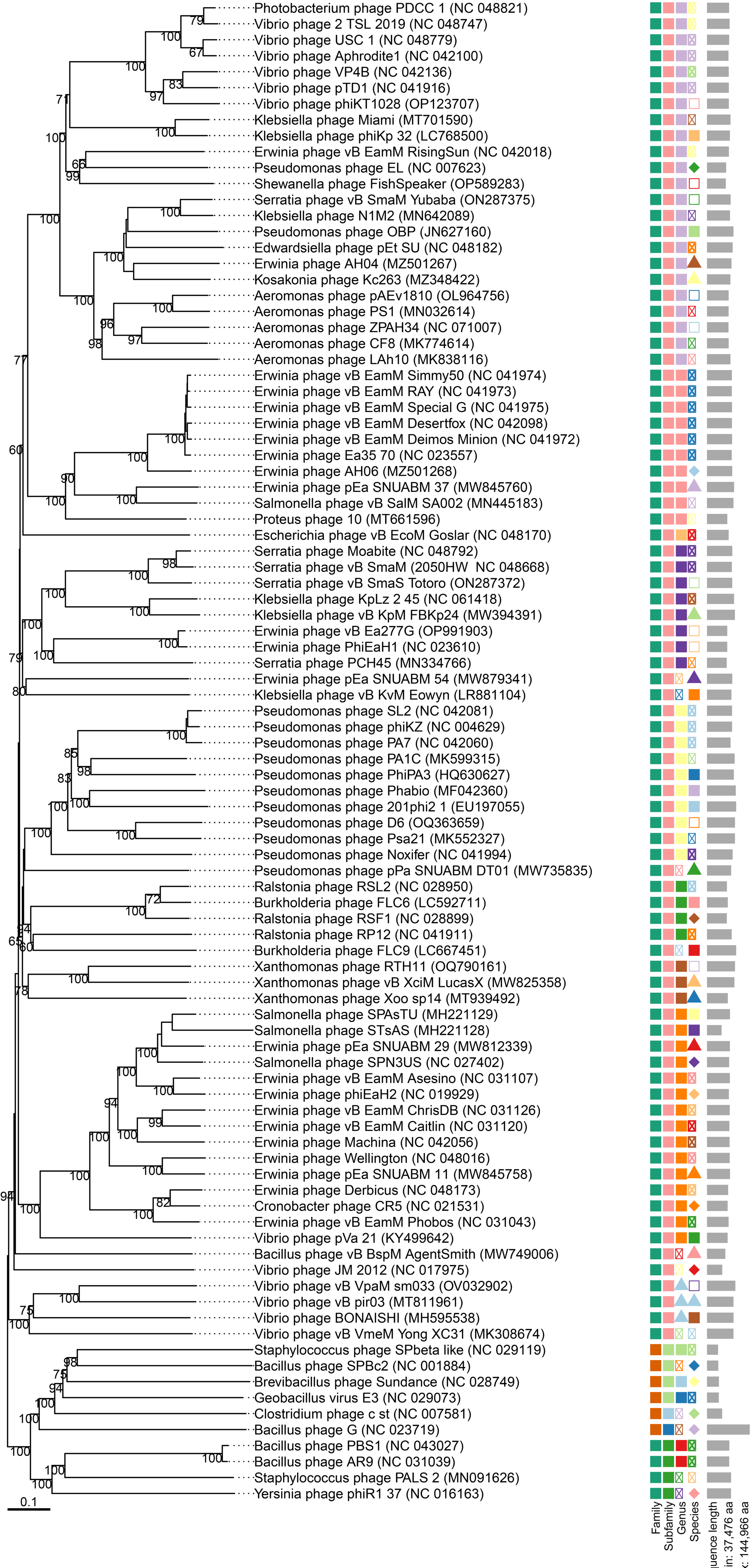


VICTOR aa tree with suggested taxa (trimming, D6)
